# A hypothesis for theta rhythm frequency control in CA1 microcircuits

**DOI:** 10.1101/2020.12.23.424154

**Authors:** Frances K Skinner, Scott Rich, Anton R Lunyov, Jeremie Lefebvre, Alexandra P Chatzikalymniou

## Abstract

Computational models of neural circuits with varying levels of biophysical detail have been generated in pursuit of an underlying mechanism explaining the ubiquitous hippocampal theta rhythm. However, within the theta rhythm are at least two types with distinct frequencies associated with different behavioural states, an aspect that must be considered in pursuit of these mechanistic explanations. Here, using our previously developed excitatory-inhibitory network models that generate theta rhythms, we investigate the robustness of theta generation to intrinsic neuronal variability by building a database of heterogeneous excitatory cells and implementing them in our microcircuit model. We specifically investigate the impact of three key ‘building block’ features of the excitatory cell model that underlie our model design: these cells’ rheobase, their capacity for post-inhibitory rebound, and their spike-frequency adaptation. We show that theta rhythms at various frequencies can arise dependent upon the combination of these building block features, and we find that the speed of these oscillations are dependent upon the excitatory cells’ response to inhibitory drive, as encapsulated by their phase response curves. Taken together, these findings support a hypothesis for theta frequency control that includes two aspects: (i) an internal mechanism that stems from the building block features of excitatory cell dynamics; (ii) an external mechanism that we describe as ‘inhibition-based tuning’ of excitatory cell firing. We propose that these mechanisms control theta rhythm frequencies and underlie their robustness.

## 1 INTRODUCTION

Hippocampal theta rhythms (≈ 3-12 Hz) observed in local field potential (LFP) recordings are associated with cognitive processes of memory formation and spatial navigation (Colgin, 2013, 2016; Hinman et al., 2018). Exactly how theta rhythms emerge is a complicated and multi-layered problem, but it is known that there are two types, denoted type 1 and type 2, that have high (7-12 Hz) or low (4-7 Hz) frequencies respectively. Type 2, but not type 1, rhythms are dependent on cholinergic drive (Bland, 1986; Buzsáki, 2002; Kramis et al., 1975). In rodents, it has been shown that social stimuli elicit high theta, and fearful stimuli elicit low theta (Tendler and Wagner, 2015), and type 2 theta oscillations have been shown to be associated with increased risk-taking behaviour (Mikulovic et al., 2018). In humans, theta frequencies are lower overall (Jacobs, 2014), but it is still possible to distinguish high and low theta frequencies, with low theta supporting encoding and retrieval of memories (Kota et al., 2020). Clearly, theta frequency control is functionally important.

It is now well-documented that theta rhythms can be generated intra-hippocampally, emerging spontaneously from an isolated whole hippocampus preparation *in vitro* (Goutagny et al., 2009). Simultaneous access to cellular and population output presents an opportunity to untangle cellular and population dynamics of how theta rhythms are generated. In previous work, we took advantage of this and built cellular and microcircuit models that could generate theta rhythms with parameters directly constrained by experimental data from the whole hippocampus preparation and the experimental literature (Ferguson et al., 2013, 2015a, 2017). Motivated by the perspective presented by Gjorgjieva et al. (2016), we considered a ‘building blocks for circuit dynamics’ analysis approach in our microcircuit model design (Ferguson et al., 2017). In this perspective, biologically known cellular, synaptic and connectivity characteristics are considered as building blocks for circuit dynamics. For example, one such cellular ‘building block’ is post-inhibitory rebound (PIR), which has previously been invoked as a contributor to the generation of cortical oscillations (McCormick et al., 2015).

In this paper we use our theta-generating microcircuit model to develop a hypothesis of how the theta frequencies could be controlled. We first describe the model microcircuit design and then assess the robustness of theta generation in the model by considering heterogeneous pyramidal (PYR) cell populations. From this, we use phase response curves (PRCs) and show that inhibitory inputs affect the theta frequency. We thus propose a hypothesis for theta frequency control in CA1 microcircuits that is dependent on internal features of PYR cells and ‘inhibition-based tuning’ of PYR cell firing. We summarize our study in schematic form in **Fig** 1.

**Figure 1.**
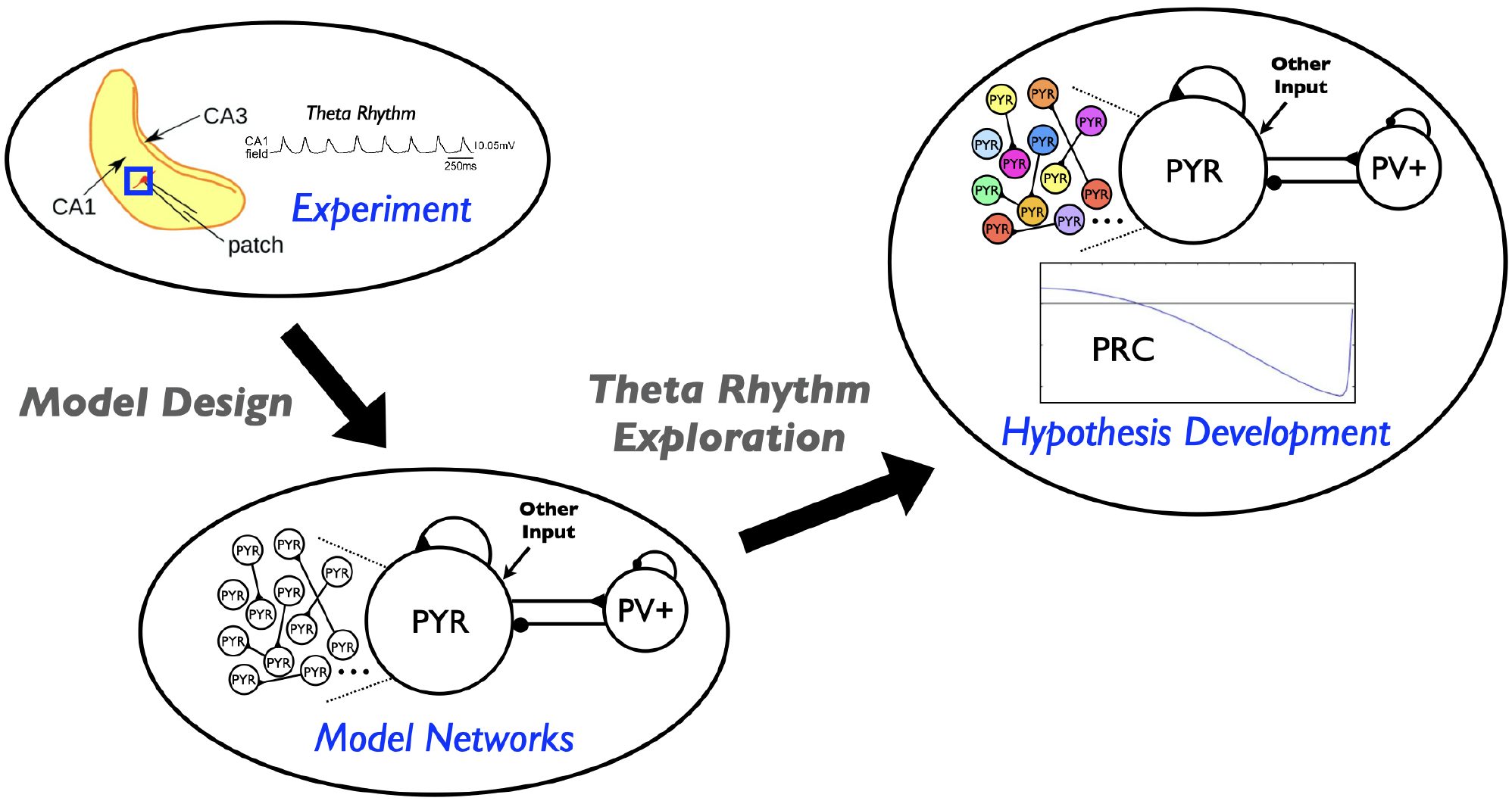
Schematic showing aspects involved in the hypothesis developed in this study. Theta rhythms are generated intrinsically in a whole hippocampus preparation of Goutagny et al. (2009) (‘Experiment’). Their generation is captured in a microcircuit model design by Ferguson et al. (2017) (‘Model Networks’). In the present paper we assess the robustness of this model design and develop a hypothesis for theta frequency control (‘Hypothesis Development’).

## 2 A DESIGN OF MICROCIRCUIT MODELS THAT PRODUCE THETA RHYTHMS

We have built cellular-based excitatory-inhibitory (E-I) network models (Ferguson et al., 2017) to understand how the intrinsic theta rhythms observed in a whole hippocampus preparation by Goutagny et al. (2009) could be generated. The model networks (see **Fig** 1 schematic) are designed to represent a ‘piece’ of the CA1 region of the hippocampus - approximately one mm^3^ that was determined to be enough to self-generate theta rhythms. It includes only two distinct cell types, pyramidal (PYR) cells and fast-firing parvalbumin-positive (PV+) cells, as represented by a single compartment model with an Izhikevich mathematical model structure (Izhikevich, 2006). The model network consists of 10,500 cells (10,000 PYR cells and 500 fast-firing PV+ cells) (Ferguson et al., 2013, 2015b). We note that we have taken advantage of a scaling relationship between cell number, connection probability and excitatory synaptic weight that allowed us to use 10,000 PYR cells rather than the 30,000 cell number size as estimated for the ‘piece’ of tissue.

We examined our models from a ‘building block for circuit dynamics’ perspective (Gjorgjieva et al., 2016) to determine if theta rhythms (i.e., theta frequency population bursts) could be generated according to experimental constraints. We first found that experimentally constrained PYR cell network models (E-cell networks alone) could generate population bursts of theta frequency (Ferguson et al., 2015b), suggesting that a cellular ‘building block’ feature of spike frequency adaptation (SFA) present in the constrained PYR cell models could be an important contributor to theta rhythm generation. However, we also found that in these E-cell only networks the PYR cells do not fire sparsely as was observed experimentally (Huh et al., 2016). When we included PV+ cells to create E-I model networks, population bursts of theta frequency were still possible and were now associated with sparse PYR cell firing in accordance with the experimental data. As the addition of PV+ cells allows PIR to be possible in the PYR cells, we consider PIR as another building block feature of importance in generating these intrinsic theta rhythms. Along with SFA and PIR features, the PYR neurons have an inherent rheobase (Rheo) feature, which is the amount of current required to make the PYR cell spike (derived from fitting to the experimental data in Ferguson et al. (2015a)). We consider this to be a third building block feature for theta rhythm generation. Further, for the model output to be consistent with experimental observations of excitatory postsynaptic current (EPSC) and inhibitory postsynaptic current (IPSC) amplitude ratios, we found that the connection probability from PV+ to PYR cells was required to be larger than from PYR to PV+ cells - a particular prediction that has been examined and found to be consistent with empirically derived connectivities (Chatzikalymniou et al., 2020).

## 3 AN ASSESSMENT OF THE MODEL DESIGN FOR ROBUST THETA RHYTHMS

In our previous work, we did not specifically examine the sensitivity of theta rhythms to SFA, PIR or Rheo features. To address this here, we create a model database of 10,000 PYR cell models. While there are various ways in which a model database could be created, we do this by simply varying specific parameter values of the PYR cell model in a regular fashion. The PYR cell model parameter values determined from fits to the experimental data (Ferguson et al., 2015a) are considered as ‘default’ values. Details for the model database creation are provided in the Appendix of the Supplementary Material.

From the created model database of PYR cell models, we obtain varied SFA, PIR and Rheo features. We define SFA, PIR and Rheo feature quantifications in the following fashion: the larger the quantified SFA value is, the stronger is the amount of the PYR cell adaptation, i.e., we get more reduction in the PYR cell spike frequency for a fixed amount of input current; the more negative the quantified PIR value is, the larger is the hyperpolarizing step required to generate a spike at the end of the step; the larger the quantified Rheo value is, the more input is required to cause the cell to spike. Details are provided in the Appendix of the Supplementary Material. For the PYR cell model with default parameter values as used in Ferguson et al. (2017), the quantified values for the building block features are: SFA = 0.46 Hz/pA, Rheo = 4.0 pA, and PIR = −5.0 pA. We refer to these as ‘base’ values. Here, with a created database of PYR cell models, we obtain a range of building block feature values distributed as shown in **Fig** 2. Further details are provided in the Appendix of the Supplementary Material.

**Figure 2.**
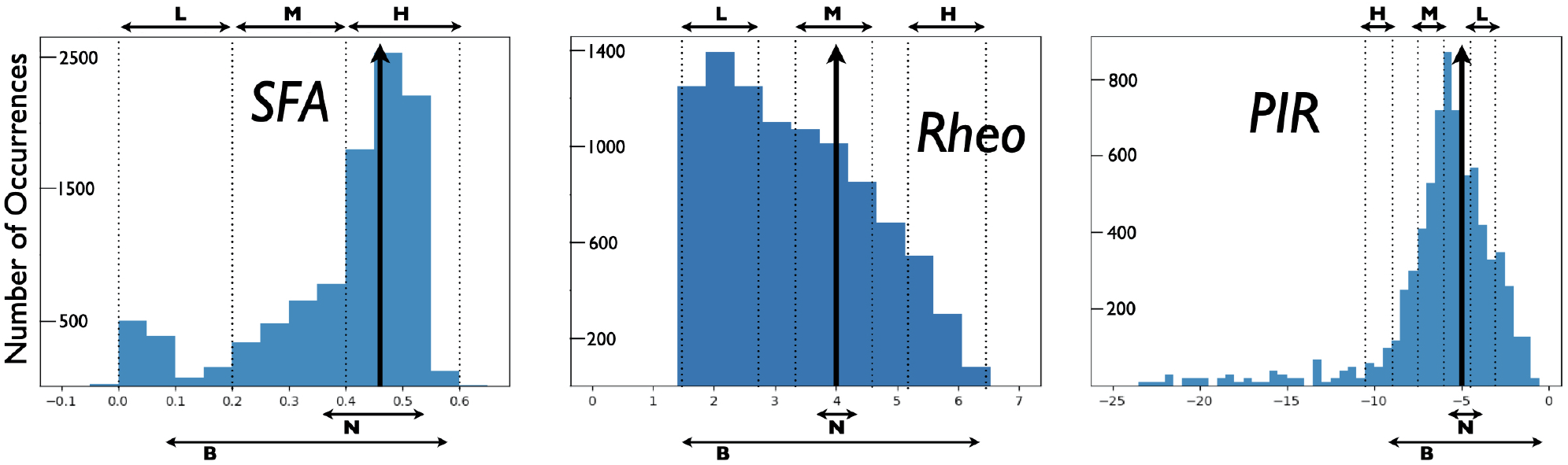
Distributions of PYR cell features from created model database. A heterogeneous set of PYR cells was created and their ‘building block’ features of SFA, Rheo and PIR were quantified. Details of this quantification are provided in the Appendix of the Supplementary Material. Histograms show the number of occurrences of SFA [=] Hz/pA, Rheo [=] pA, PIR [=] pA values, and vertical black arrows indicate [SFA,Rheo,PIR] base values. Also shown are narrow (N) and broad (B) subsets of heterogeneous PYR cell populations and low (L), medium (M) or high (H) subsets of heterogeneous PYR cell populations that do or do not include base building block values. SFA histogram has a bin resolution of 0.05, and Rheo, PIR histograms have a bin resolution of 0.5.

In the extensive E-I network simulations of Ferguson et al. (2017), the PYR cell models used were homogeneous, and all had default model parameter values. However, the networks themselves were not homogeneous because of the noisy external drives to the PYR cell models. To examine the robustness of the theta-generating mechanism in the E-I network models to variability in the SFA, PIR and Rheo features, we create heterogeneous PYR cell populations from the model database and examine whether the presence of theta rhythms in E-I networks is affected by varying these building block features.

We carry out our examination such that the heterogeneous PYR cell population in the E-I networks either does or does not include PYR cells that have base values. As a brief aside, we note that when we examine E-I networks that have homogeneous PYR cell models with parameter values different from the default ones, but that have similar SFA, PIR and Rheo base values, the resulting networks produce clear population bursts, but with a bit of variation in frequency and power. Specific examples are provided in the Appendix of the Supplementary Material.

For E-I networks with heterogeneous PYR cell populations that have PYR cells that *do* include SFA, Rheo *and* PIR base values, theta rhythms continue to be expressed. We also find that the network theta power is larger when there is a narrow rather than a broad range of values encompassing base ones. **Fig** 2 shows the narrow and broad ranges of values in our created database. Further details are provided in the Appendix of the Supplementary Material. This observation of theta power difference suggests that particular quantified feature values affect the robustness of theta rhythms since the power is larger when it more narrowly encompasses base values.

For heterogeneous E-I networks that have PYR cells that do *not* include base values for all features, we build E-I networks that have a low (L), medium (M) or high (H) range of values for SFA, Rheo and PIR features in different combinations. Thus a given heterogeneous E-I network has a triplet of [SFA,Rheo,PIR] features that have a L, M or H range of values. These values are shown in **Fig** 2. In **Fig** 3, we show the frequency (left) and power (right) of the output of these heterogeneous E-I networks designated by dots of a given color. The red circled dot is the only E-I network that *does* have base values for all of the building block features, i.e., [SFA,Rheo,PIR]=HML. We observe the following for the network frequency: Networks with Rheo=L do not produce theta rhythms when PIR and SFA= M or H; There are no theta rhythms when Rheo=M values and SFA and PIR= H; As Rheo increases, the network frequency increases, and there appears to be a stronger control of frequency by the Rheo feature relative to SFA and PIR features. For the theta power, we find that it is lowest when Rheo=L and increases as Rheo increases, but decreases as SFA or PIR increase. However, when Rheo=M, the power increases as SFA increases and as PIR decreases. From these trends, it would appear that the Rheo feature controls the theta frequency and power more than SFA or PIR. As larger values of Rheo refer to larger depolarizing currents being required for the PYR cell to fire, our observations imply that the amount of current needed for a PYR cell to fire is an essential controller of theta frequency and power, assuming that other features allow rhythms to exist in the first place. Further details from this examination are provided in the Appendix of the Supplementary Material.

**Figure 3.**
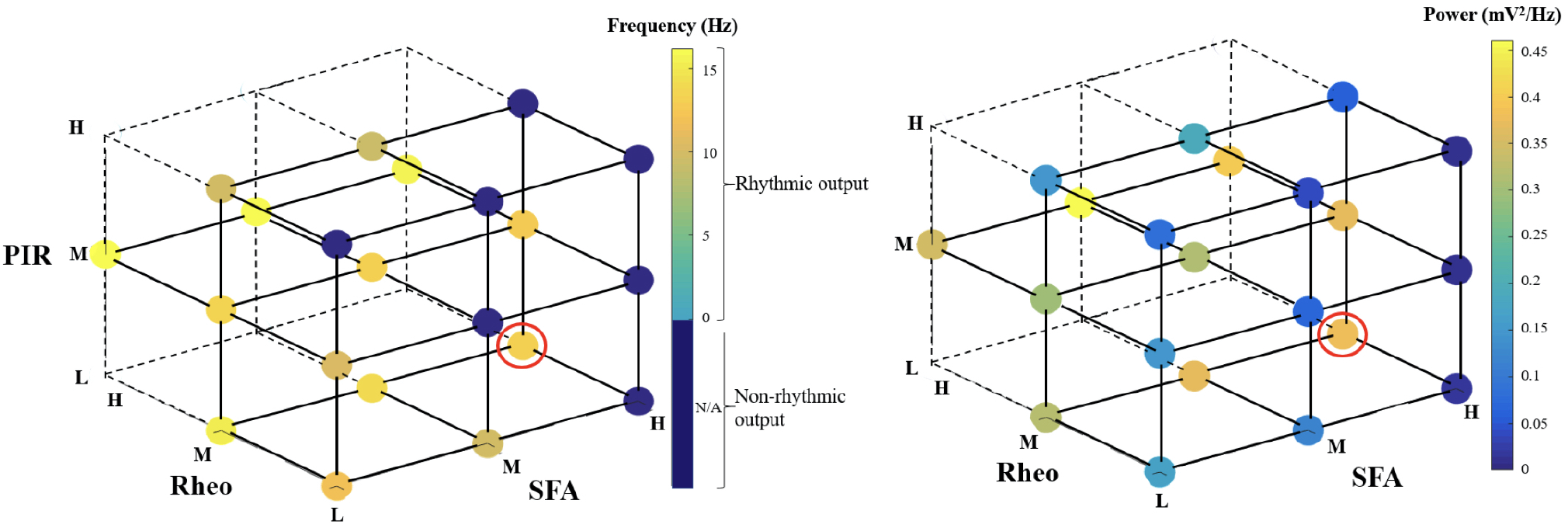
Frequency and power of theta rhythms in heterogeneous E-I networks. Each dot represents the frequency (left) or power (right) of the output of the network that has [SFA,Rheo,PIR] features with a L, M or H range of values as plotted, with the dot color representing the specific frequency or power value given in the color bar. The red circled dot is the network that has feature values that include base values for all of the features, i.e., [SFA,Rheo,PIR]=HML. The dark blue circles do not produce a rhythmic output, and the vertices that do not have any dots are where there were no individual PYR cell models to generate the particular heterogeneous network. Further details are provided in the Appendix of the Supplementary Material.

In summary, the exploration of our microcircuit model of theta rhythm generation in the whole hippocampus preparation leads us to the following conclusions regarding the influence of the three ‘building blocks’ on this dynamic: (i) a larger theta power occurs in E-I networks with heterogeneous PYR cells that include their base values and are narrowly distributed around them, and (ii) particular rheobase current values control the frequency and power of network rhythms more than the ability of the PYR cell to spike on inhibitory rebound or the particular amount of spike frequency adaptation. Thus, these simulations of E-I networks with heterogeneous PYR cell populations have allowed us to gauge the contributions of the different features and have helped us to confirm the robustness to cellular heterogeneity of the theta-generating rhythm mechanism in our microcircuit model design.

## 4 USING THE ASSESSMENT AND DESIGN TO DEVELOP A HYPOTHESIS FOR THETA FREQUENCY CONTROL

As described above, we find that large, minimally connected recurrent networks with fast-firing PV+ cells and PYR cells can produce theta frequency population rhythms consistent with experiment, driven and controlled in part by the building block features of SFA, PIR and Rheo in PYR cells. In our previous I-cell only network models of PV+ cells, coherent network output was possible with experimentally constrained PV+ cellular models and synaptic connectivities (Ferguson et al., 2013). In creating the E-I network model setup, the PV+ cell network was ‘designed’ to be in a coherent state - a function of the appropriate excitatory drive being received and the connectivity of PV+ cells. Specifically, we chose the synaptic weight (between PV+ cells) to be such that it could be at the ‘edge’ of firing coherently (high frequency) or not (see Fig. 3 in Ferguson et al. (2013)), and as such, given an appropriate excitatory drive from the PYR cells, the PV+ cell network could be in a high frequency coherent regime and be considered to be producing an inhibitory ‘bolus’ to the PYR cells. This is an important consideration for our phase response curve (PRC) considerations below.

From the several model sets of heterogeneous E-I model network outputs described in the previous section, we choose three that exhibit strong population rhythms of different frequencies. Details on these three chosen networks (specifically the heterogeneous PYR population as well as the classification of their rhythms as ‘strong’) can be found in the Appendix of the Supplementary Material. Raster plot outputs of the PYR cells in these chosen heterogeneous E-I networks are shown in **Fig** 4 where the different rhythms are referred to as ‘slow’, ‘medium’ and ‘fast’. Given the minimal nature of the microcircuit model, the frequencies of these rhythms fall a bit outside theta ranges (higher) for some networks, although the underlying theta generation mechanism and the model design is the same.

**Figure 4.**
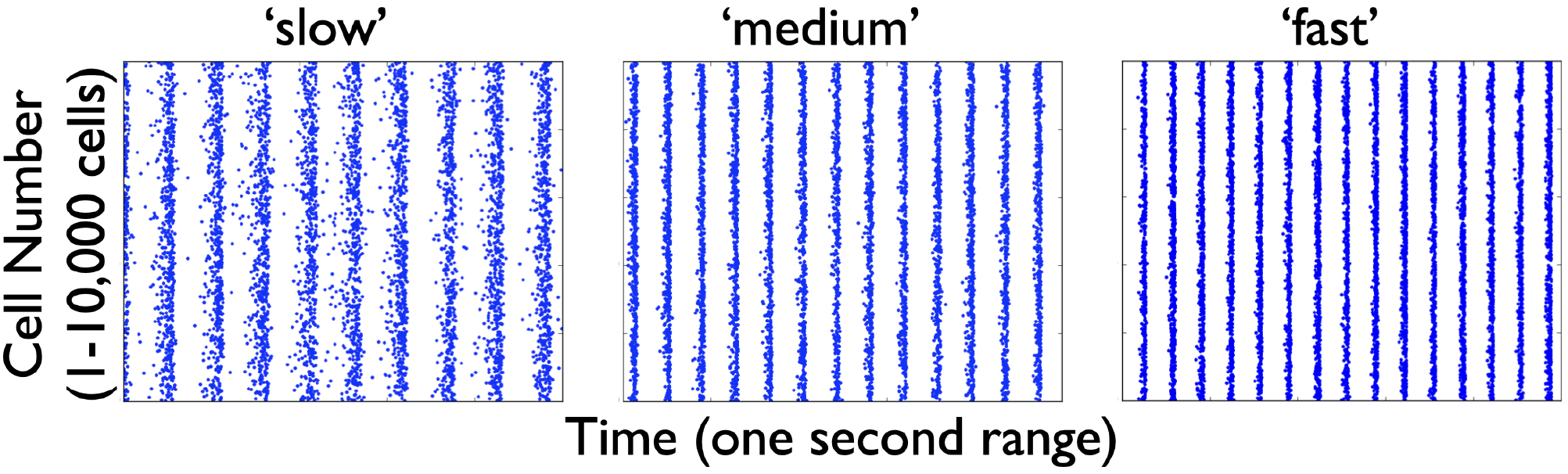
Raster plot outputs of PYR cells from three heterogeneous E-I simulations. These three model sets generating population burst rhythmic output exhibit three different frequencies that we refer to as ‘slow’ (9.6 Hz), ‘medium’ (13 Hz) and ‘fast’ (15 Hz) from their respective model sets. For all three sets, the heterogeneous PYR cells include those with Rheo base values, whereas only the model set producing the ‘medium’ output has PYR cells with SFA base values. Except for the model set producing ‘slow’ output, PYR cells have PIR base values. That is, the triplet [SFA,Rheo,PIR] feature for the slow, medium and fast networks are MMH, HML and LML respectively.

Let us now take advantage of our microcircuit design to examine how these frequencies are controlled by turning to PRC considerations (Schultheiss et al., 2011). We note that while PRCs are commonly calculated using a brief, strong, excitatory current pulse as a perturbation, we slightly modify that paradigm here and intead use a negative pulse whose amplitude and duration is motivated by the type of synaptic inputs generated during an ‘inhibitory bolus’ in our network model (see **Fig** 5). We know that the PYR cell network can generate theta population bursts on its own given its cellular adaptation characteristics (SFA feature) (Ferguson et al., 2015b). While on their own the PYR cells do not fire sparsely as in experiment, they do when a PV+ cell population is included (Ferguson et al., 2017). We consider that the resulting frequency of the E-I network’s population bursts is due to a combination of the individual PYR cell’s firing frequency and how much an inhibitory input could advance or delay the PYR cell spiking (as quantified by PRCs). The setup to consider this is schematized in **Fig** 5 and consists of the following: Each PYR cell in the heterogeneous population receives excitatory input from other PYR cells as well as a noisy drive (other input). The amount of input a PYR cell receives would of course fluctuate over time, but under reasonable approximation the PYR cell receives a mean excitatory input of about 20 to 30 pA. This approximation is based on the fact that in our E-I network models (see **Fig** 1), theta population bursts occur when PYR cells receive a zero mean excitatory drive with fluctuations of ≈ 10-30 pA (Ferguson et al., 2017). We then calculate PRCs as described above. The inhibitory pulse can advance or delay the subsequent PYR cell’s spike as quantified by the PRC, which in turn is dependent on the PYR cell’s intrinsic properties. All of these aspects are schematized in **Fig** 5.

**Figure 5.**
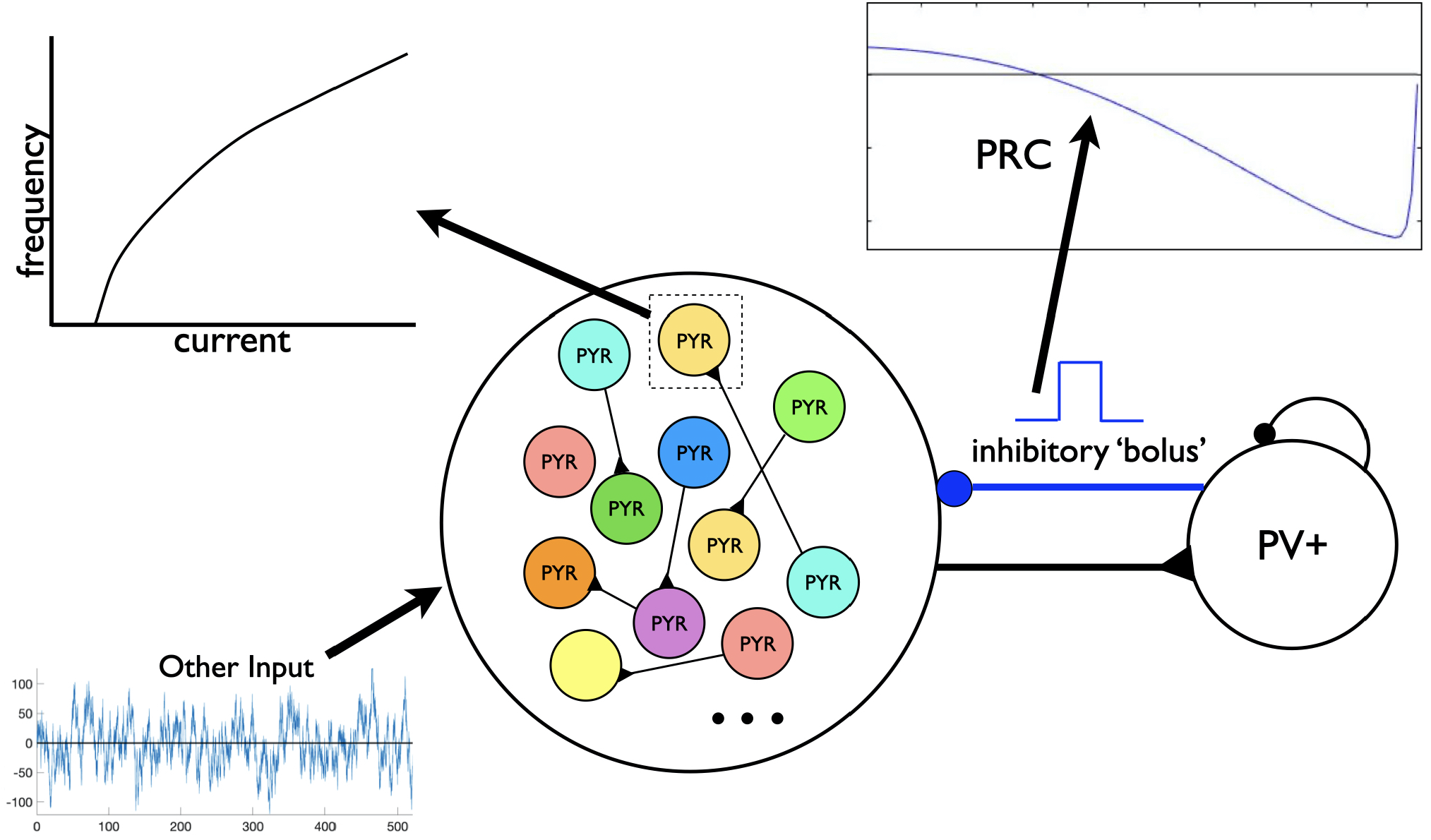
Schematic of setup for phase response curve (PRC) calculations. Assuming a theta-generating mechanism based on model design, PRCs are generated based on an inhibitory input (’bolus’) coming from the PV+ cell network to a PYR cell in the PYR cell network. Each PYR cell is receiving a noisy drive shown as ‘Other Input’, and an illustrative f-I curve is shown for one of the PYR cells. An illustration of a computed PRC based on the inhibitory input to a particular PYR cell is also shown. It would be dependent on the particular PYR cell’s model parameter values that dictates its f-I curve.

We consider the three cases of heterogeneous E-I networks exhibiting different population burst frequencies shown in **Fig** 4 and described as having a ‘slow’, ‘medium’ or ‘fast’ population burst frequency output. We generate PRCs for the several PYR cell models in the population for each of these model sets that produce the different frequency population burst outputs. Each PYR cell model in the heterogeneous population has particular PRC characteristics due to its given model parameter values, and thus exhibits a specific intrinsic frequency for a given input.

### 4.1 PRC calculations

These proceed as follows: A set input current (20:2:30 pA) is tonically applied to the model cell, and the period (defined *λ*) of the cell’s firing is calculated as the time between the ninth and 10th cell spike. The inverse of the period represents the firing frequency of the cell, reported as averages and standard deviations for entire model sets. We compute the phase response of a model neuron to a perturbation at 100 equidistant times in its normal firing cycle, where the perturbation is a 1 ms current pulse with −500 pA amplitude (as mentioned previously, considered an approximation of the synaptic input received by these cells following an ‘inhibitory bolus’). For 1 ≤ *i* ≤ 100, we define 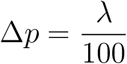 and deliver the perturbation at *i* * Δ*p* ms after the 10th cell spike. We then measure the time between the 10th and 11th cell spike as the “perturbed period” (defined *λ_p_*). We calculate the difference between this and the previously calculated period (in the absence of any perturbation) and normalize this by the normal firing period, meaning that in the PRC plots the y-axis is 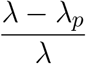. This means that negative values plotted in the PRC correspond with a phase-delay, i.e. the perturbed period was longer than the unperturbed period, and vice-versa. The x-axis in the PRC plots are the normalized time at which the perturbation was delivered, simply calculated as 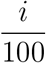. We note that we perform this calculation separately for each *i*, i.e. we re-initialize the cell and let it respond naturally to a tonic input until the 10th spike for each value of *i*, rather than perform these perturbations sequentially and risk confounding the responses.

In **Fig** 6**B** and **C** we quantify aspects of the PRC curves. In **Fig** 6**B** we simply extract the value of the normalized phase difference from the mean PRC curve for a perturbation delivered at a normalized phase of 0.3 (denoted by the arrows overlaid on **Fig** 6**A**). In **Fig** 6**C**, we quantify one aspect of the mean PRC curve’s rate of change, specifically the variability of the difference quotient calculated at each phase step, in the following straightforward way: first, this difference quotient is calculated for all but the last value of the normalized phase; second, the variance of these data is calculated simply using the *var* function in MATLAB.

**Figure 6.**
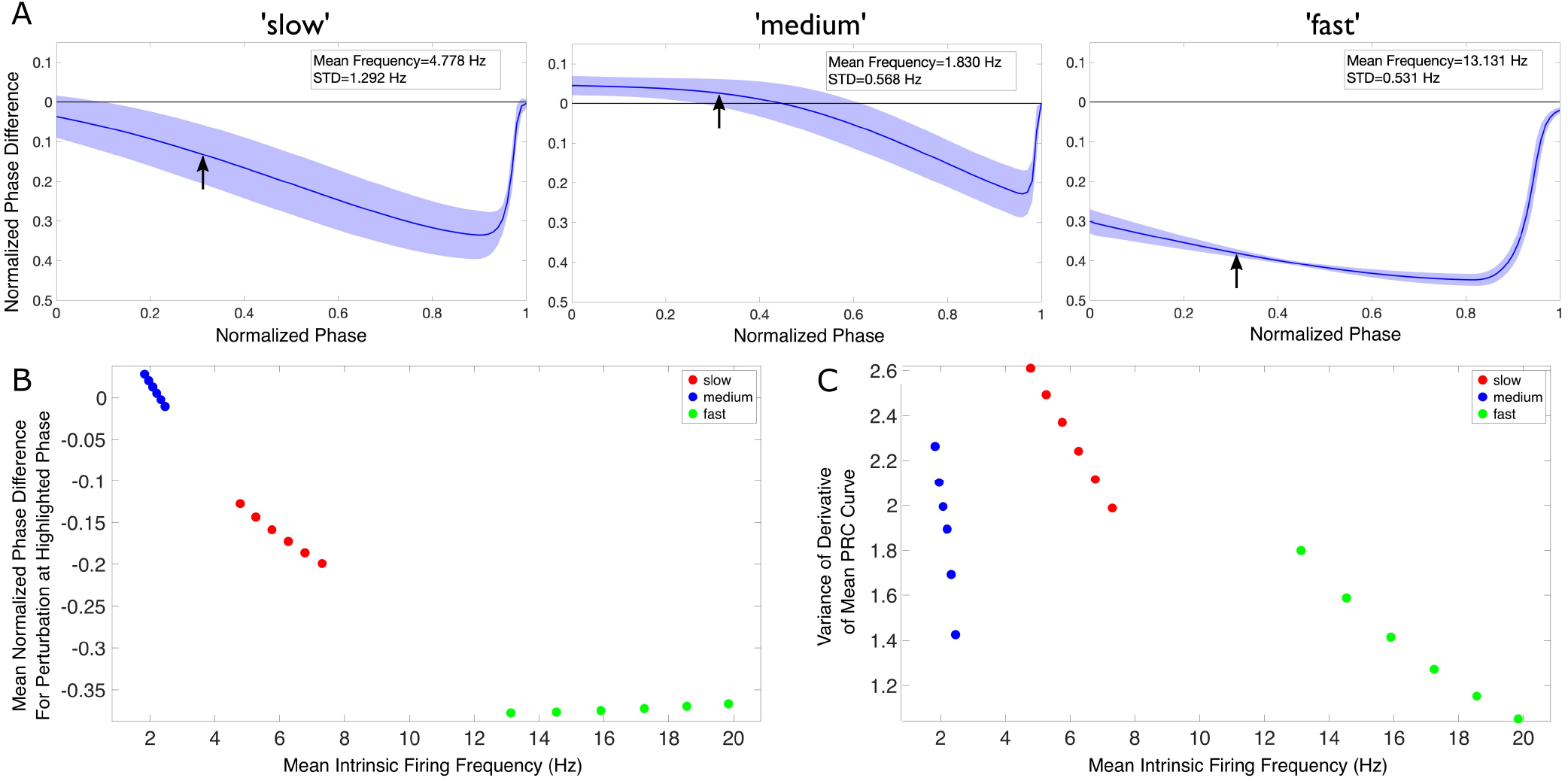
Theta rhythm frequency is influenced by inhibitory drive as quantified via PRCs and firing frequencies of individual PYR cells. **(A)** Mean PRC (solid line) for the heterogeneous PYR cell population involved in ‘slow’ (left), ‘medium’ (middle), and ‘fast’ (right) theta oscillations, calculated with an input current of 20 pA and with the shading representing ± the standard deviation. There are 25, 556 and 74 different PYR cell models in the 10,000 PYR cell populations of slow, medium and fast cases respectively. More details are provided in the Appendix of the Supplementary Material. The mean and standard deviation of the firing frequencies of the PYR cells at this input level are included in the inset of each panel. **(B-C)** After calculating both the mean PRC and mean intrinsic firing frequency for the PYR cell populations associated with our ‘slow’ (red), ‘medium’ (blue), and ‘fast’ (green) theta oscillations for six input currents (20:2:30 pA), we extract a particular feature of the mean PRC (the mean phase shift caused by a perturbation delivered at a phase of 0.3 in panel **B** and the variance of the mean PRC’s derivative in panel **C**) and plot it against the mean intrinsic firing frequency. In neither case is there a linear relationship between either axis and the theta rhythm frequency, indicating that it is a more complex combination that determines the population frequency. Note that, given the monotonic relationship between the input current and firing frequency in this range, the leftmost point for each color represents an input current of 20 pA, with each subsequent point moving rightwards representing the next input current step.

The code for generating and plotting these PRCs can be found at https://github.com/sbrich/Theta_PRCs. PRCs for input currents other than 20 pA that is shown in **Fig** 6**A** can be found at https://osf.io/yrkfv/).

### 4.2 Observations

In **Fig** 6 we first show an example of PRCs calculated for an input current of 20 pA (**Fig** 6**A**). PRCs are calculated for each model in a particular model set of heterogeneous PYR cell models, with the averaged curve presented along with a range of ± one standard deviation (shown by the shading around the curve in each plot of **Fig** 6**A**). These PRCs showcase distinct features: for instance, the PYR cells in the medium case uniquely exhibit a region of phase-advance, while the PYR cells in the fast case have the largest phase delay for perturbations delivered at all but the latest phases. Clear distinctions between the PRCs for each model set persist for all the input currents used.

To better visualize the influence of the intrinsic properties of the PYR neurons on theta rhythm frequency, we plot an extracted feature of the mean PRC against the mean firing frequency of these model sets for each of our computed input currents in **Fig** 6**B** and **C**, with the corresponding theta rhythm frequencies associated with each model set denoted by the data point’s color, with the extracted PRC features in each case described in the previous section. These visualizations clearly illustrate that *both* the PRC and the mean intrinsic firing frequency of the PYR neurons in a given model set contribute to the overall theta rhythm frequency; otherwise, these points would be “flat” with respect to either the x or y axis. Furthermore, the relationship between the extracted PRC feature of interest and the mean intrinsic firing frequency varies notably depending on the output theta rhythm frequency: for instance, in **Fig** 6**B** both the ‘slow’ and ‘medium’ model sets show a monotonically decreasing relationship between the extracted PRC value and the mean intrinsic firing frequency, while the ‘fast’ model set shows a monotonically increasing relationship. Taken together, these results show that it is a combination of the inhibitory drive and the PYR cell’s excitability that contributes to the overall theta rhythm frequency.

The intrinsic properties quantified by the PRCs help articulate potential mechanisms by which these differing theta rhythm frequencies arise. For instance, while the PYR cells in the fast case have the fastest individual firing frequencies (notably faster than what is seen in population models), their PRCs may be illustrative of how the inhibitory ‘bolus’ decreases this firing frequency towards the theta range. Meanwhile, the PYR cells in the medium case have the slowest individual firing frequencies, although they participate in ‘medium’ theta rhythm frequencies. The PRC in this case, particularly the region of phase-advance, may elucidate how inhibitory synaptic input actually accelerates PYR cell activity. These particular examples rely upon the PRC feature extracted and plotted in **Fig** 6**B**.

This analysis of the PRC features of our model sets supports our hypothesis that the frequency of the network population bursts are due to a combination of the inputs that the PYR cells receive and the intrinsic properties of those cells dictating their responses to said inputs. The cells’ response to excitatory drive is quantified in part by the mean intrinsic firing frequency of the model sets, while their response to inhibitory drive is quantified by the properties of the computed PRCs. However, this is all in the context of being able to have a stable population burst in the first place, as given by our model design with SFA, PIR and Rheo features: our models include a PYR cell population that can generate theta frequency population bursts on its own, with the PV+ cell population serving to facilitate sparse PYR cell firing. The PRC calculations here show that an appropriate inhibitory input contributes to the resulting population burst frequency.

## 5 DISCUSSION

Several models of theta rhythms have been developed (Ferguson and Skinner, 2018; Kopell et al., 2010), but they have not specifically looked at theta frequency control as coupled with its generation in an experimental context. Here, we have used a microcircuit model, as designed to generate theta rhythms representing those observed in a whole hippocampus preparation, to develop a hypothesis for theta frequency control. Our work has allowed us to propose a hypothesis for theta frequency rhythm control that encompasses two aspects: (i) an internal mechanism that stems from SFA, PIR and Rheo building block features of PYR cells; (ii) an external mechanism that involves an ‘inhibition-based tuning’ of PYR cell firing. From our previous work we already knew that minimally connected PYR cell networks produced theta frequency population bursts on their own (Ferguson et al., 2015b), but the majority of the PYR cells would fire during population theta bursts which is unlike the experimental observations of sparse PYR cell firing. With the inclusion of PV+ cells to create E-I networks, the population of PYR cells fired sparsely in accordance with experiment. It makes sense that the addition of inhibitory cells leads to less firing of PYR cells due to potential silencing from the inhibition. That theta rhythms of strong power can still emerge despite the participation of fewer PYR cells in the rhythm is likely due to the PV+ cells tuning the otherwise diverse frequencies of the PYR cells to similar frequencies, enabling this smaller group of cells to produce strong rhythms. This constitutes a main part of our proposed hypothesis. Relatedly, it has been shown that feedforward inhibition plays a role in maintaining low levels of correlated variability of spiking activity (Middleton et al., 2012).

It is important to highlight two key aspects that underlie our proposed hypothesis. First, the PYR cell population needs to be large enough so that it can collectively generate a strong excitatory drive to the inhibitory PV+ cells, and in turn the PV+ cell population should be able to fire enough (and coherently) to create a strong inhibitory ‘bolus’ to tune the PYR cell population output. Second, the net input (recurrent excitation, excitatory drive, incoming inhibition) received by the PYR cells leads to the generation of theta rhythms and its resultant frequency. It is interesting to note that similarities exist between these key aspects and the “PING mechanism” underlying the generation of gamma rhythms in E-I networks (Kopell et al., 2010; ter Wal and Tiesinga, 2013), especially considering recent research showing that rhythms with frequencies approaching the theta range can arise in PING-motivated networks (Rich et al., 2017).

We do not know whether a clear relationship between PYR cell inputs and network frequency as described in the second key aspect above actually exists, and it would be highly challenging to directly examine this experimentally. However, it is possible to use detailed, biophysical network models to explore this and gain biological insights. We have done this by bringing together the described microcircuit model used herein and a detailed, full-scale CA1 microcircuit model (Bezaire et al., 2016), and examining how the theta network frequency produced by the detailed model depends on the net input received by the PYR cells (Chatzikalymniou et al., 2020). We found that the biologically detailed models strongly support this dependence and thus our proposed hypothesis for theta rhythm frequency control. Thus, this indicates that theta frequencies in the biological system may be controlled in such a fashion.

In the previous work of Ferguson et al. (2015a), we had created PYR cell models that were either strongly adapting based on fits to the experimental data, or weakly adapting based on another experimental dataset. In Ferguson et al. (2015b), when either PYR cell models were used in E-cell only networks, that could produce theta frequency population bursts. As discussed in Ferguson et al. (2015a), it is unlikely that there are distinct types of biological PYR cells that are strongly or weakly adapting, but rather a continuum of adaptation amount dependent on the underlying balances of biophysical ion channel currents. Our explorations of the robustness of the theta generation mechanism in the microcircuit model here revealed that the frequency and power of theta rhythms were not strongly controlled by SFA feature values relative to Rheo feature values. Thus, although we created the model database starting from the strongly adapting PYR cell model parameter basis, it likely would not have mattered if the robustness examination of theta rhythm generation had been undertaken using weakly adapting PYR cell models instead.

It is perhaps not surprising that Rheo feature values are the main controller of the existence of theta rhythms and their frequency and power, as the particular Rheo value dictates whether a PYR cell would spike or not. We note that the experimental findings of Goutagny et al. (2009) had already suggested the importance of PIR in the generation of theta rhythms. In actual CA1 PYR cells, it has been shown that PIR spiking does occur, mediated by h-channels, and is locally controlled by biophysical ion channel balances (Ascoli et al., 2010). Whether PYR cells actually fire due to PIR during ongoing theta rhythms may or may not be the case, and one could potentially disentangle this in the models. However, the hypothesis developed in this work points to a confluence of features that culminate in the net current to individual PYR cells being a focus of theta rhythm frequency control. Thus, changes in the net drive to PYR cells or changes to the PYR cell’s intrinsic properties such as h-currents that would affect PIR would be expected to affect the resulting theta rhythm frequency.

PRC theory has been used in a variety of ways in the Neuroscience field (Schultheiss et al., 2011), and particularly in consideration of network dynamics. For example, Hansel et al. (1995) used PRCs to explain the differential capacity for excitatory drive to synchronize networks of Type I or Type II neurons (these types are differentiated by their bifurcation type (Izhikevich, 2006)), Rich et al. (2016) analyzed synchronization features in purely inhibitory networks using PRCs, and Achuthan and Canavier (2009) used PRCs to understand clustering in networks. We took advantage of PRC theory by considering phase-resetting of the PYR cells due to incoming inhibitory input. In this way, we were able to hypothesize an inhibition-based tuning mechanism for control of the theta rhythm frequency based on the PRC shape (amount of advance or delay) and the PYR cell’s intrinsic firing frequency. Our use of PRCs relied on our observations of the effect of different PRC shapes on the resulting theta rhythm. For example, such a consideration was used by Rich et al. (2016) to explain differential synchrony patterns in inhibitory networks of Type 1 vs Type II neurons.

In conclusion, we have developed a hypothesis for how theta rhythm frequencies are controlled in the CA1 hippocampus. This hypothesis is built on the theta-generating mechanism of the microcircuit model design. Even though it does not include all of the known inhibitory cell types, it perhaps captures essential elements in play in biological circuits and may apply more widely in the brain regarding the generation and control of theta rhythm frequencies.

## AUTHOR CONTRIBUTIONS

FS, JL, AC contributed to conception and supervision of the study. SR, AL performed computations and analyses. FS wrote the first draft of the manuscript. SR wrote sections of the manuscript. All authors contributed to manuscript revision, read, and approved the submitted version.

## ACKNOWLEDGMENTS

Parts of this work have been released as a preprint (Chatzikalymniou et al., 2020).

## SUPPLEMENTAL DATA

There is a supplementary file referred to as the Appendix in the main text.

## DATA AVAILABILITY STATEMENT

Additional datasets generated by this study that are not present here or in the Appendix (Supplementary Material) can be found at https://osf.io/yrkfv/.

