## Supplementary Appendix for "A hypothesis for theta rhythm frequency control in CA1 microcircuits"

### Supplementary Material

#### APPENDIX

In this Appendix we provide details of:

- mathematical equations and parameter values
- pyramidal (PYR) cell model database setup and feature quantifications
- excitatory-inhibitory (E-I) network simulation setup and output specifics

##### Cellular specifics and equations

The network structure and cellular details for the model simulations in the present paper are as in Ferguson et al. (2017). That is, cellular models of excitatory, pyramidal (PYR) and inhibitory, parvalbumin-positive (PV+), fast-firing cells are based on experimental data from the *in vitro* whole hippocampus preparation (Ferguson et al., 2013, 2015). They use the mathematical model structure developed by Izhikevich (Izhikevich, 2010, 2006), in which the subthreshold behaviour and the upstroke of the action potential are captured, and a reset mechanism to represent the spike's fast downstroke is used. Despite being relatively simple, parameter choices can be made such that they have a well-defined (albeit limited) relationship to the electrophysiological recordings. It has a fast variable representing the membrane potential, $V\ (mV)$, and a variable for the slow "recovery" current, $u\ (pA)$. We used a slight modification to be able to reproduce the spike width. It is described by the following set of equations:

$$\begin{aligned} C_m\dot{V} &= k(V - v_r)(V - v_t) - u + I_{other} - I_{syn} \\ \dot{u} &= a[b(V - v_r) - u] \end{aligned} \tag{S1}$$

$$\text{if } V \geq v_{peak},\ \text{then } V \leftarrow c,\ u \leftarrow u + d$$
$$\text{where } k = k_{low} \text{ if } V \leq v_t,\ k = k_{high} \text{ if } V > v_t$$

where $C_m\ (pF)$ is the membrane capacitance, $v_r\ (mV)$ is the resting membrane potential, $v_t\ (mV)$ is the instantaneous threshold potential, $v_{peak}\ (mV)$ is the spike cut-off value, $a\ (ms^{-1})$ is the recovery time constant of the adaptation current, $b\ (nS)$ describes the sensitivity of the adaptation current to subthreshold fluctuations - greater values couple $V$ and $u$ more strongly resulting in possible subthreshold oscillations and low-threshold spiking dynamics, $c\ (mV)$ is the voltage reset value, $d\ (pA)$ is the total amount of outward minus inward currents activated during the spike and affecting the after-spike behaviour, and $k\ (nS/mV)$ represents a scaling factor. $I_{syn}$ = 0 for the isolated cell. $I_{other}$ is as described below for computing metrics for the PYR cell or E-cell.

Model parameter values for the PV+ cell or I-cell (units above) are: $v_r$=-60.6; $v_t$=-43.1; $v_{peak}$=-2.5; $c$=-67; $k_{high}$=14; $C_m$=90; $a$=0.1; $b$=-0.1; $d$=0.1; $k_{low}$=1.7. These parameters are as previously determined

(Ferguson et al., 2013), and are not varied. Model parameter values (units above) for the PYR cell are: $v_r$=-61.8; $v_t$=-57; $v_{peak}$=22.6; $c$=-65.8; $k_{high}$=3.3; $C_m$=115; $a$=0.0012; $b$=3; $d$=10; $k_{low}$=0.1. These parameters are as previously determined for strongly adapting cells (Ferguson et al., 2015). We refer to them as default parameters, and specifically, the $a, b, d, k_{low}$ parameters are varied.

#### Network specifics and equations

The cellular models described above were used to create excitatory-inhibitory (E-I) networks as done in Ferguson et al. (2017). Specifically, synaptic input within the PYR cell (E-cell) PV+ cell (I-cell) populations, and between PYR and PV+ cells, is represented in Equation S1 as:

$$I_{syn} = g \cdot s(V - E_{rev}) \tag{S2}$$

where $g\ (nS)$ is the maximal synaptic conductance of the synapse from a presynaptic neuron to the postsynaptic neuron, $E_{rev}\ (mV)$ is the reversal potential of the synapse, and $V\ (mV)$ is the membrane potential of the postsynaptic cell. The gating variable, $s$, represents the fraction of open synaptic channels, and is given by first order kinetics (Destexhe et al. (1994), and see p.159 in Ermentrout and Terman (2010)):

$$\dot{s} = \alpha[T](1 - s) - \beta s \tag{S3}$$

The parameters $\alpha$ (in $mM^{-1}ms^{-1}$) and $\beta$ (in $ms^{-1}$) in Equation S3 are related to the inverse of the rise and decay time constants ($\tau_R,\ \tau_D$ in $ms$). $[T]$ represents the concentration of transmitter released by a presynaptic spike. Suppose that the time of a spike is $t = t_0$ and $[T]$ is given by a square pulse of height $1\ mM$ lasting for $1\ ms$ (until $t_1$). Then, we can represent

$$s(t - t_0) = s_\infty + (s(t_0) - s_\infty)e^{-\frac{t-t_0}{\tau_s}}, \quad t_0 < t < t_1$$

where $s_\infty = \frac{\alpha}{\alpha+\beta}$ and $\tau_s = \frac{1}{\alpha+\beta}$. After the pulse of transmitter has gone, $s(t)$ decays as

$$s(t) = s(t_1)e^{-\beta(t-t_1)}$$

For network simulations, $I_{other}$ in Equation S1 represents 'other input' to the PYR cell population is given by $I_{other} = -g_e(t)(V - E_{rev})$. $g_e(t)$ is a stochastic process similar to the Ornstein-Uhlenbeck process as used by Destexhe and colleagues (Destexhe et al., 2001)

$$\frac{dg_e(t)}{dt} = -\frac{1}{\tau_e}(g_e(t) - g_{e,mean}) + \sqrt{\frac{2\sigma_e^2}{\tau_e}}\chi_e(t) \tag{S4}$$

where $\chi_e(t)$ is an independent Gaussian white noise process of unit standard deviation and zero mean, $g_{e,mean}\ (nS)$ is the average conductance, $\sigma_e\ (nS)$ is the noise standard deviation value, and $\tau_e$ is the time constant for excitatory synapses. $\tau_e$ is fixed based on values as used in Destexhe et al. (2001) ($\tau_e = 2.73\ ms$).

Parameter values (rationale and references given in Ferguson et al. (2017)) are: $E_{rev}$= -15 or -85 mV for excitatory or inhibitory reversal potentials respectively. Rise and decay time constants are, respectively, 0.27 and 1.7 msec for PV+ to PV+ cells; 0.3 and 3.5 msec for PV+ to PYR cells; 0.37 and 2.1 msec for PYR to PV+ cells; 0.5 and 3 msec for PYR to PYR cells. Connection probabilities are fixed at 0.12 for

PV+ to PV+ cells and 0.01 for PYR to PYR cells, as estimated from the literature. For the simulations in this paper, we use connection probabilities that were found to be in line with the experimental data. That is, the connection probability from PV+ to PYR cells ($c_{PV,PYR}$) is larger than from PYR to PV+ cells ($c_{PYR,PV}$).

For the heterogeneous networks examined in this paper, we mainly focus on parameter values from Table 5 of Ferguson et al. (2017): $g_{pyr}$=0.094 nS, $\sigma_e$=0.6 nS, $g_{pyr-pv}$=3 nS, $g_{pv-pyr}$=8.7 nS, $c_{PYR,PV}$=0.02, $c_{PV,PYR}$=0.3, $g_{e,mean} = 0$ nS. An actual instantiation of the 'other input' that these parameter values produce can be seen in the schematic figure in the main text (**Figure 4**). We also consider networks with parameter values of: $g_{pyr}$=0.014 nS, $\sigma_e$=0.6 nS, $c_{PYR,PV}$=0.02, $c_{PV,PYR}$=0.3; and $g_{pyr}$=0.084 nS, $\sigma_e$=0.2 nS, $c_{PYR,PV}$=0.04, $c_{PV,PYR}$=0.5; and $g_{pyr}$=0.084 nS, $\sigma_e$=0.6 nS, $c_{PYR,PV}$=0.02, $c_{PV,PYR}$=0.5 ($g_{pyr-pv}$, $g_{pv-pyr}$, $g_{e,mean}$ the same as focused parameter values), and similar results are obtained. From the model we know that theta population bursts occur when PYR cells receive zero mean excitatory drive with fluctuations of $\approx$ 10-30 pA (as estimated from 0.2 to 0.6 nS 'noise') (Ferguson et al., 2017).

#### PYR cell (E-cell) model database and quantifying building block features

To create a database of PYR cell models, we range the $a, b, d, k_{low}$ model parameter values to create 10,000 models, using 10 different values for each parameter such that they encompass the default values of: $a$=0.0012$ms^{-1}$; $b$=3.0$nS$, $d$=10$pA$, $k_{low}$=0.10$nS/mV$. The parameter values range from zero to about twice the default values such that the default values are about the midpoint values. Specifically, they are (*[initial value, final value, resolution]*): $a$ = [0.0, 0.00216, 0.00024]; $b$ = [0.0, 5.4, 0.6]; $d$ = [0, 18, 2]; $k_{low}$ = [0.0, 0.18, 0.02].

For each PYR cell model, spike frequency adaptation (SFA), post-inhibitory rebound (PIR) and rheobase (Rheo) features are quantified to allow comparisons to be made. Quantification of features is done as follows:
*Rheo:* Starting from $v_r$, each PYR cell model is given a constant current from -25 to 25 pA in 0.5 pA increments. If a spike is generated within the first 500 msec, then that constant current value is considered as the rheobase current, and is taken as the *Rheo* quantified value.
*PIR:* Starting from $v_r$, each PYR cell model is subjected to a one second hyperpolarizing step current for current values from 0 to -25 pA with a resolution of 0.5 pA. If a spike occurred upon termination of a given hyperpolarization step (i.e., a PIR spike) but not at the previous step value, then that step value is considered as the *PIR* quantified value.
*SFA:* Starting from $v_r$, each PYR cell model is subjected to input currents for one second, from 0 to 98 pA (inclusive) in 2 pA increments. For each input current, the number of spikes is recorded, and the interspike interval is calculated between the first and second spikes, and the last and second from last spike. The inverse is taken and defined as the initial and final frequency at that current. The initial and final frequencies as a function of the current steps creates a smooth, approximately linear relationship, so lines are fitted to the initial and final frequency plots. The slopes of those lines are subtracted from one another (the initial slope is always steeper) to produce the *SFA* quantified value.

The range of quantified values obtained from the model database of 10,000 PYR cells is: SFA: -0.001 to 0.64 (Hz/pA); Rheo: 1.5 to 6.5 (pA); PIR: -23.5 to -1.0 (pA). How they end up being distributed is shown in **Figure 2** of the main text, and while clearly not a uniform or normal distribution, they encompass a range of values which is what we seek. A PIR spike was not obtained within the hyperpolarizing step range explored for 2,570 of the 10,000 PYR cells in the model database, and were thus excluded from the PYR cell models to be used in the E-I network simulations. The quantified values for the strongly adapting PYR cell model (see above for full model and parameter values) are: SFA= 0.46; Rheo= 4.0; PIR= -5.0. We refer to them as the base values.

#### Heterogeneous PYR cell groupings

The two ways in which heterogeneous PYR cell populations are created for use in the E-I network simulations is as follows:

(i) Using narrow (*N*) or broad (*B*) ranges of values for [SFA, Rheo, PIR] relative to base values, where *N* or *B* means that [SFA, Rheo, PIR] metric values are $\pm$ [0.1, 0.5, 0.5] or $\pm$ [0.45, 3.0, 5.0] respectively, of base values. Thus, *NNN* refers to models with [SFA, Rheo, PIR] values of: [(0.36 to 0.56) exclusive of bounds; 4.0; -5.0], and *BBB* refers to models with [SFA, Rheo, PIR] values of: [(0.01 to 0.64) exclusive of bounds (noting that 0.64 is the maximum possible in the model database set); 1.5, 2.0, 2.5, 3.0, 3.5, 4.0, 4.5, 5.0, 5.5, 6.0, 6.5; -0.5, -1.0, -1.5, -2.0, -2.5, -3.0, -3.5, -4.0, -4.5, -5.0, -5.5, -6.0, -6.5, -7.0, -7.5, -8.0, -8.5, -9.0, -9.5]. Note that since the resolution of the Rheo and PIR quantified values are 0.5, and the manner in which it is defined (see above), the *N* range for Rheo has models in which Rheo = 4.0 only, and similarly, the *N* range for PIR has models in which PIR = -5.0 only.
The other sets (using ranges as defined above) have quantified values as follows: *BBN*=[(0.01 to 0.64) exclusive of bounds; 1.5, 2.0, 2.5, 3.0, 3.5, 4.0, 4.5, 5.0, 5.5, 6.0, 6.5; -5.0]; *BNB*=[(0.01 to 0.64) exclusive of bounds; 4.0; -0.5, -1.0, -1.5, -2.0, -2.5, -3.0, -3.5, -4.0, -4.5, -5.0, -5.5, -6.0, -6.5, -7.0, -7.5, -8.0, -8.5, -9.0, -9.5]; *NBN*=[(0.36 to 0.56) exclusive of bounds; 1.5, 2.0, 2.5, 3.0, 3.5, 4.0, 4.5, 5.0, 5.5, 6.0, 6.5; -5.0]; and so on for *BNN*, *NBB*, and *NNB*. These eight possible cases and the number of models in each of them is given in **Table** S2, along with the population frequency and power. Parameter value histograms for each of these combinations from the model database set are given in `https://osf.io/yrkfv/`, and what ranges of the quantified values in the database that they encompass is shown in the main text (**Figure 2**).

(ii) Using low (*L*), medium (*M*) or high (*H*) values, with SFA quantified value ranges exclusive of endpoints given as: SFA: *L* = [(0.0 to 0.2)], *M* = [(0.2 to 0.4)], *H* = [(0.4 to 0.6)]; Rheo: *L* = [1.5, 2.0, 2.5], *M* = [3.5, 4.0, 4.5], *H* = [5.5, 6.0, 6.5]. PIR: *L* = [-3.5, -4.0, -4.5], *M* = [-6.5, -7.0, -7.5], *H* = [-9.5, -10.0, -10.5]. This means that the base values fall into the *HML* case, with the small caveat that the PIR base value is just outside the *L* range. The gaps in these ranges are due to the automation of the exploration and to ensure that there is no overlap in the quantified values for a given case. Note that there ended up being no models for the cases: *HHH, HHL, MHH, MHL, LHH, LHL*, from the created model database. Thus there are 21 cases from the generated model database, and the number of models present in each case is given in **Table** S2. Parameter value histograms for 12 of these cases are given in `https://osf.io/yrkfv/`, and what ranges of the quantified values in the database that they encompass is shown in the main text (**Figure 2**).

#### E-I network setup and simulations

To build E-I model networks, we choose PYR cells from the model database in two ways in consideration of SFA, Rheo and PIR features, referring to them as a trio in the following order: [SFA, Rheo, PIR]. The chosen PYR cells are distributed among the 10,000 cells to be used in the E-I network simulations in the following way: An individual PYR cell model is randomly chosen from the set of models of a particular heterogeneous PYR cell population that have [SFA, Rheo, PIR] values within the specified range. For example, if there are 33 PYR cell models in the set, then the number of cells conforming to each of the 33 PYR cell models should approach 10,000/33 in the E-I network, but there may not be an exactly equal number of the different PYR cell models. That is, we do the following: If there are 33 PYR cell models in the given heterogeneous PYR cell model set, then each PYR cell model out of 10,000 in the E-I network is given a random number between 1 and 33, and assigned that model's parameters. We note that comparisons between the heterogeneous E-I networks are not perfectly ideal since the number of different PYR cell models varies (see **Table** S2), and so the 'amount' of heterogeneity would vary in the various E-I networks. However, since we are mainly considering whether the theta rhythm would be lost or not, this is deemed to be acceptable.

The model E-I network simulations are done using the Neuroscience Gateway (NSG) for high-performance computing (Sivagnanam et al., 2013). Simulations are run for 10 seconds using the Euler integration method to integrate the cell equations with a timestep of 0.04 msec, and 2nd order Runge-Kutta with a timestep of 0.04 msec for network simulations. The frequency and network power of the network simulation is computed as before (Ferguson et al., 2017). That is, for each network simulation, the population activity is defined as the average membrane potential of all the cells, with the frequency and network power taken as frequency and spectral peak from a fast Fourier transform (FFT) calculation of the population activity.

Code details are provided in `https://github.com/FKSkinnerLab/CA1_Minimal_Model_Hetero` and simulation output in `https://osf.io/yrkfv/`.

#### E-I network simulation output details

In the extensive E-I network simulations of Ferguson et al. (2017), the PYR cell models used were homogeneous, all having default model parameter values. However, the networks themselves were not homogeneous because of the noisy external drives to the PYR cell models. To examine the robustness of the theta-generating mechanism in the E-I network models with consideration of the SFA, Rheo and PIR features, we create heterogeneous PYR cell populations from the model database that has a range of feature values to examine whether the presence of theta rhythms (population bursts of theta frequency) is dependent on particular feature values.

We carry out our examination such that the heterogeneous PYR cell population in the E-I networks either do or do not include PYR cells that have base values. When we examined E-I networks of homogeneous PYR cell models with parameter values different from the default ones, but with similar values for quantified features, the resulting networks produce clear population bursts. Specific examples are shown in **Table** S1 along with their model parameter and quantified feature values. The fact that the rhythm is not lost in any of these networks with homogeneous model parameter values already suggests that the populations bursts are not particularly sensitive to the specific SFA quantified values as the rhythm isn't lost as SFA varies. However SFA has some effect on the specific power and frequency of the population bursts.

We classify the PYR cells from the created model database to build E-I model networks with heterogeneous PYR cell populations in two groups according to their quantified values of the [SFA, Rheo, PIR] feature trio. The first group corresponds to: (i) Narrow ($N$) or broad ($B$) ranges of [SFA, Rheo, PIR] values that include the base values, and the second group corresponds to: (ii) Low ($L$), medium ($M$) or high ($H$) ranges of [SFA, Rheo, PIR] values that do not necessarily include the base values. These groups are shown in the main text paper (**Figure 2**).

For each group we create networks corresponding to combinations of the quantified values of the SFA, Rheo, PIR feature ranges. For (i), there are eight possible E-I network cases from $N$ and $B$ combination sets and the number of models in each case is given in **Table** S2, along with the frequency and power of

**Table S1.** E-I Network Simulation Examples with Homogeneous PYR cell models.

| ***Homogeneous cells in network*** Model ID # | ***Parameter values*** $(a, b, d, k_{low})$ *Units:* (1/ms, nS, pA, nS/mV) | ***Quantified values*** *(SFA, Rheo, PIR)* *Units:* (Hz/pA, pA, pA) | ***Power*** ($mV^2$/Hz) | ***Frequency*** (Hz) |
|---|---|---|---|---|
| Original (base) | (0.0012, 3.0, 10, 0.10) | (0.46, 4.0, -5.0) | 0.36 | 12.2 |
| # 7 | (0.00072, 3.6, 18, 0.16) | (0.51, 4.0, -5.0) | 0.21 | 11.8 |
| # 32 | (0.00072, 4.8, 12, 0.16) | (0.51, 4.0, -5.0) | 0.37 | 14.2 |
| # 56 | (0.00096, 3.6, 4, 0.12) | (0.38, 4.0, -5.0) | 0.40 | 13.6 |
| # 81 | (0.00096, 4.2, 12, 0.10) | (0.49, 4.0, -5.0) | 0.42 | 13.8 |
| # 115 | (0.0012, 3.6, 14, 0.06) | (0.49, 4.0, -5.0) | 0.34 | 13.0 |

the particular network. For (ii), there are 27 possible network cases from *L, M* and *H* combination sets and the number of models in each case is also given in **Table** S2, along with the frequency and power of the particular network. As it turns out, there are no PYR cell models in the created model database for *HHH, HHL, MHH, MHL, LHH, LHL* network cases. We thus have simulation output for only 21 different E-I networks with heterogeneous PYR cell populations generated using (ii).

There is a clear maintenance of rhythms for the eight cases of heterogeneous group (i), as shown in the top part of **Table** S2, where the quantified values are chosen in either a narrow or broad fashion encompassing base values. Their frequencies are similar to each other and to that of the E-I network with homogeneous, default PYR cell model parameter values (see first row in **Table** S1). Interestingly, the network power is larger when there is a narrow rather than a broad range of values encompassing base values (compare *NNN* to *BBB* in **Table** S2), suggesting that particular quantified feature values affect how strong the theta frequency population bursts are, as given by their theta power. If we have heterogeneous E-I model networks with PYR cell model parameter values that have broadly distributed feature values that include base values, then population rhythms remain with less than a 1 Hz variation in population frequency. This further implies that a quantification of the features can capture the underlying E-I balances necessary for the emergence of theta frequency population bursts in the microcircuit model.

Output for the 21 cases of heterogeneous E-I networks with PYR cell models that have quantified feature values that do not necessarily encompass base values, i.e., group (ii), is shown in **Table** S2 where it is clear that a rhythm (i.e., population bursts) is not always present. We first note that the E-I network for the *HML* case is the one that mostly encompasses base values for all three features. As one might expect, the power and frequency of this E-I network case is similar to the heterogeneous (i) E-I network cases which also encompass the base values. Considering the network power values of all of these heterogeneous network cases, it is easy to see which networks are not rhythmic. Essentially, if the power is below 0.1, then there is not a clear rhythm - these cases are shown in the lower part of **Table** S2. The cases in **Table** S2 that are starred are networks that have started to lose their rhythm. To view the output from several heterogeneous E-I networks, in **Fig** S1 we show PYR cell raster plots for four cases (designated with an ‘R’ in **Table** S2). In three of them, there is still a rhythm, but there are clear frequency and PYR cell burst firing characteristic differences. In **Fig** S2 we show PYR cell raster plots for four additional cases (designated with an ‘R-supp’ in **Table** S2) for when the rhythm is lost so that the different patterning can be seen.

In considering the cases in which the rhythm is lost, it appears that the existence of the rhythm is not heavily dependent on the specific SFA quantified values, since rhythms still exist even when moving away from ”Hxx” cases (i.e., those encompassing the base SFA value) - *MML* and *LML* cases. However, the rhythm *is* lost if the E-I networks do not include base values for Rheo or PIR, specifically ”xMx” (base Rheo value) or ”xxL” (closest to base PIR value) cases. For Rheo, consider the *HLL* case (no *HHL* case

**Table S2.** Heterogeneous E-I Network Simulations.

| *Network case [SFA,Rheo,PIR]* | *Number of different PYR cell models* | *Power* ($mV^2$/Hz) | *Frequency* (Hz) |
|---|---|---|---|
| **Group (i)** | | | |
| *NNN* | 137 | 0.37 | 13.0 |
| *BBB* | 6780 | 0.27 | 13.0 |
| *BBN* | 550 | 0.28 | 12.8 |
| *BNB* | 1010 | 0.29 | 13.0 |
| *BNN* | 180 | 0.34 | 13.4 |
| *NBB* | 4955 | 0.30 | 13.0 |
| *NBN* | 416 | 0.24 | 12.2 |
| *NNB* | 729 | 0.33 | 12.6 |
| **Group (ii)** | | | |
| ***HML*** (R) | 556 | 0.38 | 13.0 |
| *HHM* | 313 | 0.40 | 15.6 |
| *HMM* | 493 | 0.37 | 12.8 |
| *MHM* | 157 | 0.46 | 15.8 |
| ***MMH*** (R) | 25 | 0.19 | 9.6 |
| *MMM* | 294 | 0.31 | 13.2 |
| *MML* (R) | 110 | 0.37 | 13.8 |
| *MLL** | 99 | 0.12 | 10.0 |
| *LHM* | 49 | 0.35 | 16.2 |
| *LMH** | 12 | 0.15 | 9.8 |
| *LMM* | 103 | 0.30 | 13.6 |
| ***LML*** | 74 | 0.32 | 15.0 |
| *LLM** | 29 | 0.15 | 10.4 |
| *LLL* | 64 | 0.17 | 12.0 |
| **No Rhythm** | | | |
| *HMH* (R-supp) | 33 | 0.06 | n/a (9.2) |
| *HLH* | 97 | 0.01 | n/a (1.2) |
| *HLM* (R) | 171 | 0.01 | n/a (0.6) |
| *HLL* | 417 | 0.02 | n/a (0.6) |
| *MLH* (R-supp) | 27 | 0.04 | n/a (8.2) |
| *MLM* (R-supp) | 50 | 0.07 | n/a (8.6) |
| *LLH* (R-supp) | 16 | 0.08 | n/a (10.0) |

Top set of eight network cases use heterogeneous PYR cell models from group (i) and the rest use heterogeneous PYR cell models from group (ii). Boldfaced cases are networks from which PRCs are explicitly used - see main paper. (R) and (R-supp) refers to networks in which PYR cell rasters from the E-I networks are explicitly shown in **Fig** S1 and **Fig** S2 respectively. (*) refers to networks that are almost losing their rhythm

to consider) and for PIR, consider the *HMH* case (less so for the *HMM* case). This allows us to suggest the following: the particular rheobase current value of the PYR cell, and the ability of the PYR cell to fire a spike with a less hyperpolarized current step are needed for the theta-generating mechanism in the microcircuit models, along with some amount of spike frequency adaptation. A detailed examination of these trends is shown in the main text (**Figure 3**).

We take advantage of these explorations to develop our hypothesis in the main text. From our E-I network simulations, we choose three strong rhythms that have different frequencies. By a strong rhythm, we mean those that have a network power greater than 0.15, as based on our examination of the outputs from all the

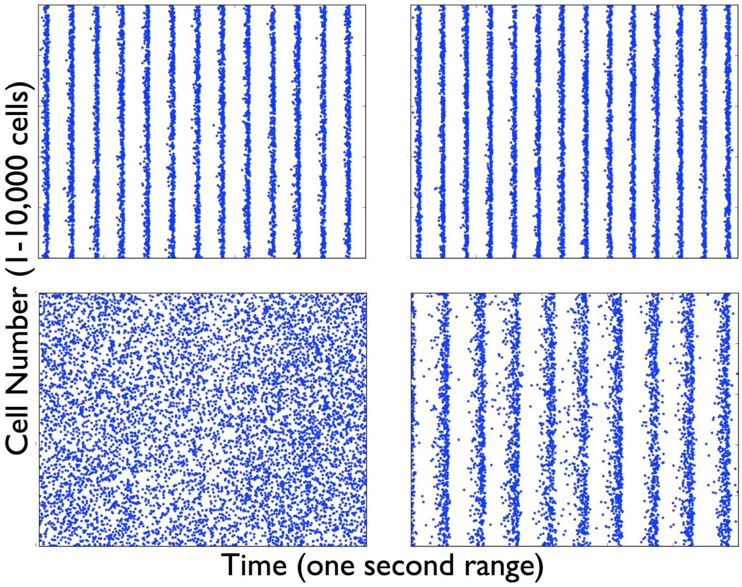


**Figure S1. Raster plots of PYR cells in heterogeneous E-I networks.**
Simulations of E-I networks with 10,000 heterogeneous PYR cells and 500 PV+ cells produce output that have PYR cell raster plots as shown here with a one second time range. The specific examples are labelled as *(R)* in **Table** S2 and refer to the following sets: HML (top-left), MML (top-right), HLM (bottom-left), MMH (bottom-right). Acronyms are defined in the main text.

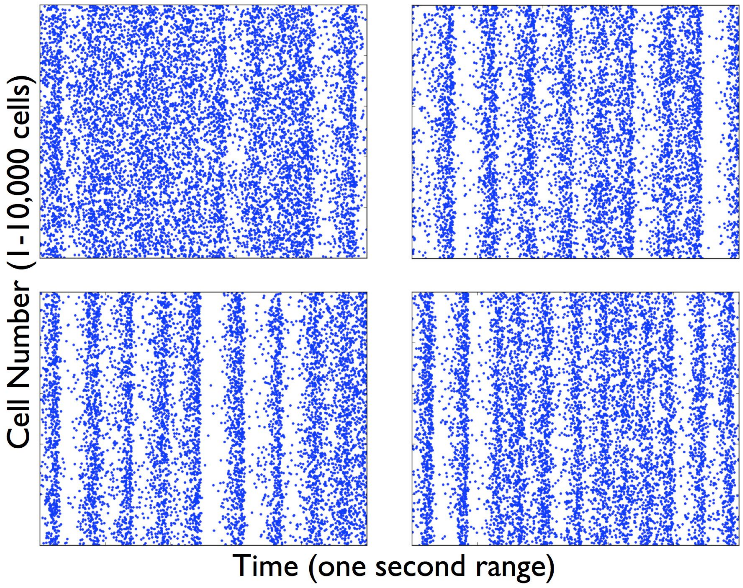


**Figure S2. Loss of Rhythm - Raster plots of PYR cells in heterogeneous E-I networks.**
Simulations of E-I networks (10,000 heterogeneous PYR cells and 500 PV+ cells with a one second time range). The specific examples are labelled as *(R-supp)* in **Table** S2 and refer to the following sets: MLH (top-left), HMH (top-right), MLM (bottom-left), LLH (bottom-right).

E-I network simulations. They are shown as boldfaced cases in **Table** S2 - HML, MMH and LML. Note that for all three of these cases, the heterogeneous PYR cell population includes Rheo base values.
